# *In silico* prediction of metabolic trait robustness in microbial cells

**DOI:** 10.64898/2026.01.09.698552

**Authors:** Chunhao Gu, Ville Mustonen, Paula Jouhten

**Affiliations:** Department of Bioproducts and Biosystems, Aalto University, Espoo, Finland; Faculty of Biological and Environmental Sciences, Organismal and Evolutionary Biology Research Programme, University of Helsinki, Helsinki, Finland

## Abstract

In industrial applications, microbial strains undergo notable biomass expansion subjecting them to Darwinian selection. The consequent adaptive evolution may threaten the often tediously developed desired traits of the strains, such as flavor or platform chemical production. Yet, it remains unresolved how to predict the evolutionary trait robustness. Here, we propose TRAEV (Trait Robustness Against EVolution), a computational framework for *in silico* prediction of evolutionary trajectories and robustness of desired metabolic traits in application environments. TRAEV uses constraint-based metabolic model simulations to predict environment-dependent trait-fitness dependencies and Monte Carlo-based perturbation analysis to account for the stochasticity of adaptive evolution. First, TRAEV predicts the immediate phenotypic adaptation to the new environment, and then, the evolutionary trajectories of fitness and desired trait by sampling enzyme usage changes. From the predicted trajectories, trait robustness is quantified as two scores: Robustness Score (RS) and Trade-off Score (TS). RS and TS are the mean of a normalized desired trait and the mean of the product of normalized changes in the desired trait and in fitness, respectively, over intermediate metabolic states along the evolutionary trajectory. We validated TRAEV by demonstrating that it predicted the relative robustness of heterologous pigmentation of genetically engineered *Saccharomyces cerevisiae* strains in synthetic defined chemical environments aligned with experimental observations. We then further applied TRAEV to predictively assess the robustness of desired aroma generation trait in a wine must environment by multiple *S. cerevisiae* strains if they were developed via a laboratory selection process. Thus, we showed how TRAEV predictions could guide such strain development. TRAEV can be integrated into computational strain design workflows across microbial strains, metabolic traits, and application environments. Ultimately, model-predicted evolutionary robustness of desired traits can guide both strain and process development and help avoiding production losses and enhancing the economic attractiveness of industrial applications using microbial cells.

**Author summary:** Microbial cells developed to express desired traits by genetic engineering or selection processes are used in a wide range of applications, from biotechnological chemical production to food and beverage fermentations. However, when the cells replicate in the application environment the traits are subject to Darwinian selection and consequent adaptive evolution. As a result, desired traits can be rapidly lost. To mitigate such undesired evolutionary changes, we developed TRAEV (Trait Robustness Against EVolution), a computational framework for *in silico* assessment of evolutionary trajectories and robustness of desired traits in application environments by integrating constraint-based modeling and Monte Carlo-based perturbation analysis methods. We demonstrate the usability of TRAEV by validating its predictions with experimental data on the robustness of heterologous pigmentation of engineered *Saccharomyces cerevisiae* strains, and applying it to predict the robustness of aroma generation in wine must by multiple *S. cerevisiae* strains if they were developed by laboratory selection in specific conditions. As being applicable across microbial strains, metabolic traits, and application environments, we believe that TRAEV can help to avoid production losses, and thus, contribute to the development of economically attractive industrial applications using microbial cells.

## Introduction

In biotechnology, microbial cells can be either genetically engineered [1, 2] or improved through artificial or natural selection [3, 4] to enhance traits of interest, most notably, the overproduction of target compounds. Such engineered or evolved microbial strains can be used in industrial production of fuels, platform chemicals [5], drugs [6], and aromas, flavors, and preservatives in food and beverages [7]. However, the enhanced traits inevitably divert the translational machinery capacity and/or metabolic precursors away from cell growth associated traits [8, 9]. This reallocation disrupts intracellular trait interdependencies and compromises cellular fitness. Thereby, the risk of evolutionary instability is elevated and a loss of the initially enhanced traits over time may occur [10].

Previous studies [11–13] have primarily focused on achieving desired metabolic traits under laboratory conditions, but often overlooked the long-term stability and evolutionary robustness of these traits in industrial application environments. However, engineered or evolved traits that allow microbial cells to achieve the desired application performance are subject to Darwinian evolution because of their prevalent fitness effects (i.e., the negative covariance between traits aligned with application objectives and cellular fitness) [14]. Consequently, trait robustness, i.e., the ability to maintain the trait under environmental (e.g., nutrient availability, pH) and genetic (e.g., mutations) perturbations and across multiple evolutionary generations, remains a fundamental challenge for large-scale industrial applications [10]. Attempts to develop strains with higher trait robustness have been made by coupling production traits to cell growth, either through the use of product sensors [15] or by reducing the metabolic network [16, 17]. Nevertheless, these approaches are limited in scope, as they rely on traits that are selectively advantageous under specific conditions. Therefore, there is a pressing need for widely applicable approaches to predict and control the resulting evolutionary trajectories, which are pivotal for producing more efficient and robust biotechnology processes [18, 19].

Over the past decade, computational modeling, particularly constraint-based modeling, has become an indispensable tool for rational strain design [20]. Genome-scale metabolic models (GEMs) are *in silico* reconstructions of cellular metabolism based on current biological knowledge, including genes, reactions, and gene–protein–reaction associations [21]. Constraint-based modeling that combines GEMs and constraint-based analysis techniques such as flux balance analysis (FBA) [22] and minimization of metabolic adjustment (MOMA) [23] has demonstrated versatile applications in metabolic engineering, including simulating flux distributions [22], identifying genetic engineering targets [24, 25], predicting metabolic bottlenecks [26], and designing selection environments [27].

Here, we developed a novel computational framework, TRAEV (Trait Robustness Against EVolution), which integrated constraint-based modeling and Monte Carlo-based perturbation analysis methods to predict relative trait robustness across strains and application environments. TRAEV uses constraint-based GEM simulations to predict an engineered or laboratory evolved strain’s immediate adjusted metabolic state due to phenotypic plasticity and the evolutionarily adapted metabolic state in an application environment. Based on these states, TRAEV estimates possible evolutionary trajectories using a Monte Carlo-based perturbation analysis method, from which the trait robustness is assessed and quantified as the Robustness Score (RS) and the

Trade-off Score (TS). We demonstrated TRAEV for predicting the robustness of heterologous pigmentation of engineered*Saccharomyces cerevisiae* strains and for screening laboratory evolution environments to develop *S. cerevisiae* strains exhibiting robust aroma generation in a wine must environment.

## Results

### TRAEV for predicting trait robustness in evolving cells

Microbial strains expressing desired traits are typically developed through genetic modification (GM), adaptive laboratory evolution (ALE), or a combination of both, under controlled laboratory conditions. However, when these strains are taken to industrial applications they become introduced into a new chemical environment (e.g., wine must or corn stalk hydrolysate media), eliciting immediate metabolic adjustments due to phenotypic plasticity [28]. This is subsequently followed over time by genotypic changes (evolutionary adaptation) driven by Darwinian evolution (Figure 1A). Both processes can alter the expression of desired traits. Therefore, an approach for predicting trait robustness in application environments is called for.

**Fig 1.**
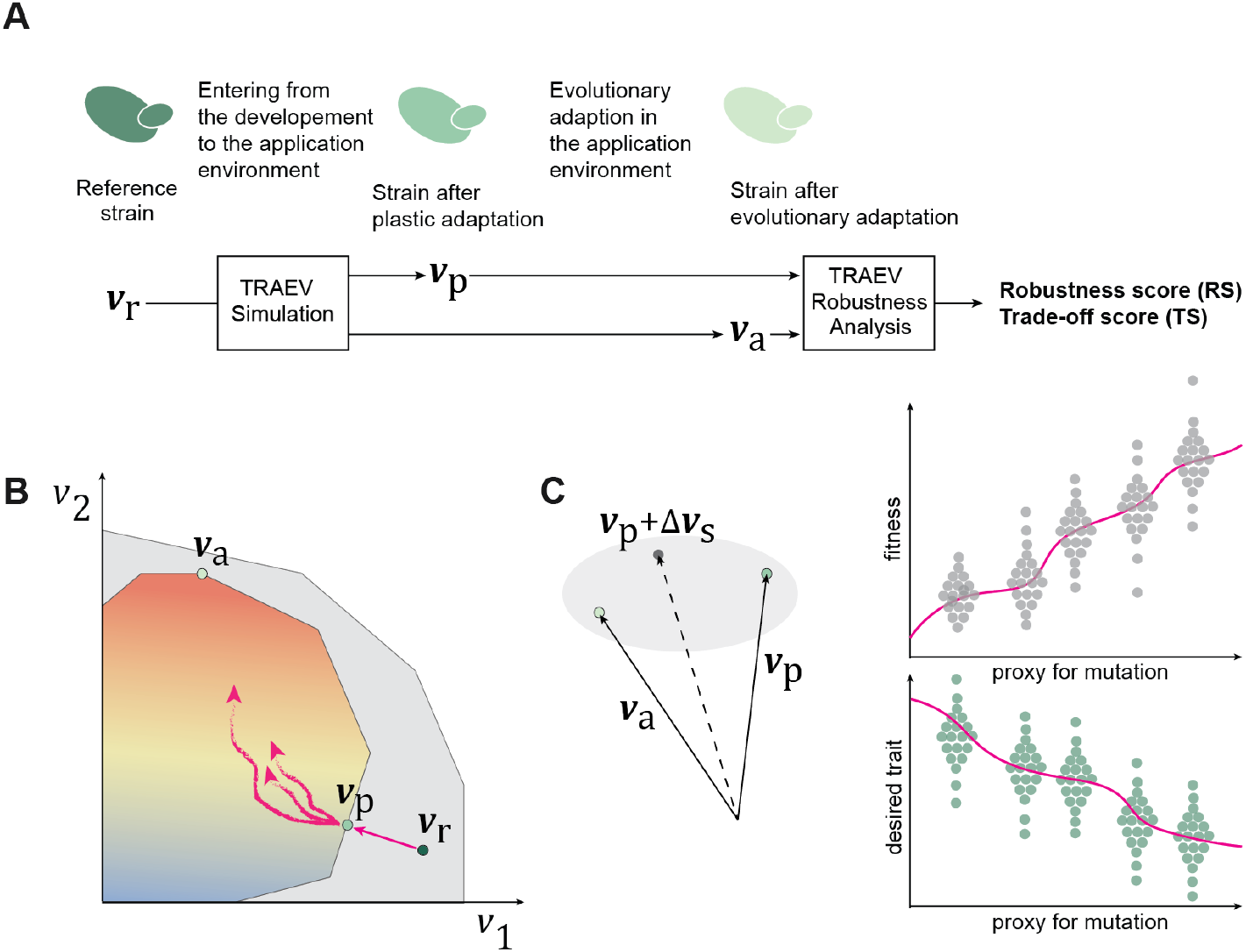
Schematic overview of TRAEV. (A) Workflow of TRAEV. The reference strain (**v**_r_) represents a strain developed through genetic modification (GM) or adaptive laboratory evolution (ALE) under laboratory conditions and computed using pFBA or SWITCHX. With **v**_r_, the TRAEV Simulation calculates the strain’s immediate phenotypically adjusted state (**v**_p_) and its evolutionarily adapted state (**v**_a_) in the application environment. Based on these states, the TRAEV Robustness Analysis estimates the evolutionary trajectories and the corresponding Robustness Score (RS) and Trade-off Score (TS). (B) Conceptual rationale of TRAEV Simulation. A two-dimensional flux space is defined by fluxes (*v*_1_, *v*_2_). Within this space, the gray polygon represents the feasible space in the development environment, within which the strain achieves the reference metabolic state **v**_r_ before being introduced into the application environment. The colored polygon represents the feasible space in the application environment, within which **v**_p_ is the state nearest to **v**_r_, while **v**_a_ is the fitness-optimal state where Darwinian evolution would on average move to. (C) Illustration of TRAEV Robustness Analysis. An evolutionarily intermediate metabolic state is represented as **v**_p_ + Δ**v**_*s*_, where Δ**v**_*s*_ = **m**_*s*_ ⊙ (**v**_a_ − **v**_p_) and **m**_*s*_ is a binary mask indicating a randomly selected subset *s*. By deriving the fitness and desired trait of these intermediate states along progressive changes in enzyme usage (serving as a proxy for mutations), TRAEV delineates the evolutionary trajectories for the calculation of RS and TS.

To this end, we developed TRAEV, a computational framework for predicting trait robustness in cells adaptively evolving in application environments. TRAEV employs constraint-based GEM simulations to predict both phenotypic plasticity and evolutionary adaptation. The simulation results are then analyzed to derive evolutionary trajectories and assess trait robustness. Specifically, TRAEV consists of two main modules: Simulation and Robustness Analysis.

As shown in Figure 1A, the Simulation takes as input a reference metabolic state (**v**_r_), which represents the metabolic state of an engineered or laboratory-evolved strain. In this study, a metabolic state is defined as a vector containing both metabolic fluxes and corresponding enzyme usages (see Methods). **v**_r_ can be computed using either parsimonious flux balance analysis (pFBA) [29] or SWITCHX (Eq. 6). Then, the Simulation calculates two key metabolic states in the application environment:

1. **v**_p_: the immediate metabolic state due to phenotypic plasticity after switching to the application environment, calculated using linear minimization of metabolic adjustment (MOMA) [30]. This state minimizes the distance from **v**_r_ while maintaining a biologically minimal growth rate to ensure viability.
2. **v**_a_: the adapted metabolic state after evolution in the application environment, calculated using SWITCHX (Eq. 6) with the adapted growth rate estimated using Algorithm 1. This state maximizes fitness while maintaining minimal changes in enzyme usage relative to **v**_r_.

A schematic illustration of the rationale of the Simulation is shown in Figure 1B.

In the Robustness Analysis, TRAEV employs a Monte Carlo-based perturbation analysis method to generate an ensemble of evolutionarily intermediate metabolic states between **v**_p_ and **v**_a_. Each intermediate state is computed using linear MOMA [30] with **v**_p_ as the reference state, while a randomly selected subset of enzyme usages is enforced to their corresponding values in **v**_a_. From the linear MOMA solution, fitness (Eq. 1) and the desired trait (Eq. 2) are calculated and then normalized (Eqs. 3, 4)). As illustrated in Figure 1C, the resulting ensemble delineates the evolutionary trajectories of fitness and the desired trait along progressive changes in enzyme usage during evolution. Finally, two scores are derived to characterize the evolution in the application environment:

1. Robustness Score (RS): a metric defined as the mean, over evolutionarily intermediate metabolic states, of the normalized desired trait, reflecting the robustness (i.e., stability and overall level) of the desired trait (Eq. 9).
2. Trade-off Score (TS): a covariance-like metric defined as the mean, over evolutionarily intermediate metabolic states, of the product of normalized changes (relative to **v**_p_) in the desired trait and in fitness, representing the trait-fitness trade-off (i.e., negative coupling) (Eq. 10).

The Robustness Score (RS) ranges from 0 to 1, with higher values indicating higher robustness of the desired trait against perturbations in metabolism. When comparing the robustness of the same desired trait across multiple strains, RS is calculated by normalizing the trait with the maximum trait value among the strains (Eq. 4), thereby representing the trajectory-averaged attainable level of the desired trait rather than stability alone. In contrast, the Trade-off Score (TS) ranges from −1 to 0, with values closer to zero indicating weaker negative coupling between the desired trait and fitness, thereby suggesting a higher potential to long-term stability of the trait. Together, RS and TS provide a quantitative framework for comparing strain performance in target application environments and for guiding rational strain design for industrial applications.

The implementation details of TRAEV are provided in the Methods section.

### Predicting trait robustness of engineered pigment production

The biosyntheses of blue indigoidine and red bikaverin are natural traits of certain bacteria (e.g., *Streptomyces* spp.) and fungi (e.g., *Fusarium* spp.), respectively. However, natural pigment synthesis is subject to complex regulation and is typically not induced under standard laboratory or industrial cultivation conditions. To overcome this limitation, heterologous production of indigoidine and bikaverin has been successfully achieved in *S. cerevisiae*, a common biotechnology host, by introducing their biosynthetic pathways via genetic engineering [11, 12].

Here, we revisited the results from a recent experiment [31], where *S. cerevisiae* CEN.PK113-7D and S288C strains were engineered for the production of indigoidine and bikaverin, respectively. Adaptive laboratory evolution (ALE) of the engineered strains were performed in two distinct media: a chemically defined medium containing ammonium as the sole nitrogen source and D-galactose as the sole carbon source (YNBG), and a rich medium containing peptone as the nitrogen source and D-galactose as the sole carbon source (YPG). In both media, after 25 transfers (i.e., ca. 175 generations in YNBG and ca. 200 generations in YPG), the bikaverin-producing colonies remained mostly pigmented, while the indigoidine-producing lineages had nearly lost all production. These results indicated that the bikaverin-producing strain exhibited higher trait robustness during prolonged cultivation (Figure 2A, B).

**Fig 2.**
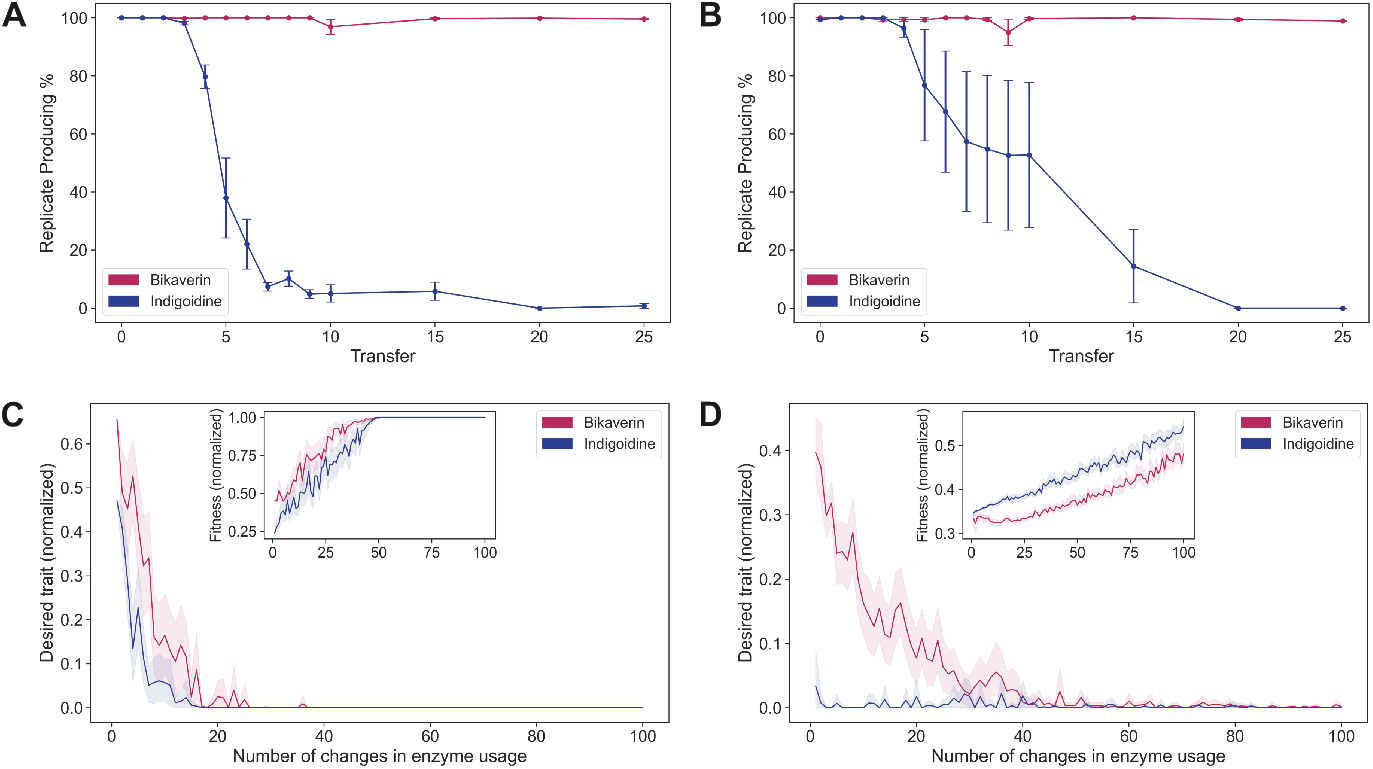
Experimental and TRAEV-predicted evolutionary trajectories of pigment-producing *S. cerevisiae* strains. (A, B) Experimental trajectories showing the percentage of pigmented colonies over 25 transfers in (A) YNBG and (B) YPG media, based on data from [31]. Each point with an error bar represents the mean (*±* SD) of four biological replicates. The bikaverin-producing strain (red) exhibited significant stability compared to the indigoidine-producing strain (blue), indicating higher trait robustness. (C, D) TRAEV-predicted trajectories illustrating changes in normalized desired trait (pigment secretion flux) vs. the number of changes in enzyme usage from 1 to 100 in (C) YNBG and (D) YPG media. Each point represents the mean of sampled metabolic states; shaded areas denote 95% confidence intervals. Insets show normalized fitness vs. enzyme usage changes. The predicted trajectories qualitatively agree with the corresponding experimental trajectories, consistently showing that the bikaverin-producing strain (red) maintains higher trait robustness than the indigoidine-producing strain (blue).

To validate whether these observations could be computationally predicted, we simulated the experiments using pFBA and TRAEV. The *S. cerevisiae* GEM [32] was augmented with the heterologous pathways for producing indigoidine [11] and bikaverin [12], respectively. pFBA was used to calculate the metabolic state **v**_r_ for each engineered strain. With **v**_r_ as the reference state of each strain, TRAEV was then performed to simulate ALE in YNBG and YPG, yielding the immediate phenotypically adjusted metabolic state (**v**_p_) and the evolutionarily adapted metabolic state (**v**_a_). Subsequently, the Robustness Analysis was applied to predict the evolutionary trajectories and calculate the corresponding Robustness Scores (Figure 2C, D). The implementation details are provided in the Methods section.

As shown in Figure 2C and D, the bikaverin-producing strains exhibited a more gradual decline in pigment production than the indigoidine-producing strains, indicating higher trait robustness. These simulated evolutionary trajectories qualitatively agreed with the experimental observations (Figure 2A, B). In the YNBG medium, the bikaverin-producing strain showed the Robustness Score (RS) of 4.61 × 10^−2^, compared to 2.12 × 10^−2^ for the indigoidine-producing strain. In the YPG medium, the difference in RS was more pronounced, with values of 5.33 × 10^−2^ for the bikaverin-producing strain and 3.22 ×10^−3^ for the indigoidine-producing strain, respectively. Together, these results showed that TRAEV could be used to predict and compare trait robustness in target environments before experimental testing.

### Model-guided design of ALE environments for adaptively evolving robust metabolic traits

In a previous study [27], a constraint-based modeling framework named EvolveX was introduced to identify selection environments for ALE that promote strains to evolve enhanced level of a desired trait once transferred to the application environment. This strategy relies on a tacking trait, a metabolic trait that is coupled to fitness (e.g., cell growth) in the selection environment and to the desired trait in the application environment. However, in the application environment, neither the tacking trait nor the desired trait is necessarily coupled to fitness. Consequently, following evolutionary adaptation driven by Darwinian selection, trait robustness in the application environment remains uncertain based on the EvolveX Scores.

Here, we used TRAEV to re-evaluate the scoring of selection environments for ALE reported in [27], aiming to obtain the robust production, in wine must environment, of two classes of aroma compounds: (1) phenylethyl alcohol and its acetate ester (PEA) and (2) branched-chain amino acid-derived higher alcohols and their esters (BCHA). We designed 1,171 distinct selection environments (see S3 File), each comprising three nutrients as carbon and nitrogen sources. For each selection environment, we used the *S. cerevisiae* GEM [32] and pFBA with nutrient and growth rate constraints to calculate the adapted metabolic state (**v**_r_) in that selection environment. With **v**_r_ as the reference state of each strain, TRAEV was then performed to simulate adaptive evolution in anaerobic wine must-like conditions with D-glucose, D-fructose, and yeast assimilable nitrogen (ammonium and free amino acids) as the carbon and nitrogen sources, obtaining the immediate phenotypically adjusted state (**v**_p_) and the evolutionarily adapted metabolic state (**v**_a_) for each strain. Finally, for both classes of aroma compounds, the Robustness Analysis was conducted to predict the evolutionary trajectories along the changes in enzyme usage and calculate the corresponding Robustness Scores and Trade-off Scores. The implementation details are provided in the Methods section.

In the TRAEV results, the strains evolved in 401 of the 1171 selection environments were able to produce PEA in the application environment, whereas the strains evolved in 1064 of these selection environments were able to produce BCHA. In Figure 3, we plotted the evolutionary trajectories of ten ALE-evolved strains for the PEA-producing trait (Figure 3A, C) and ten for the BCHA-producing trait (Figure 3B, D). The corresponding Robustness Scores (RS) and Trade-off Scores (TS) (Eqs. 9, 10) are summarized in Table 1. These examples were randomly chosen from among the ALE-evolved strains ranked in the top 5% by RS, making the trajectories representative of the robustness profiles for each trait.

**Table 1.** RS and TS of ALE-evolved *S. cerevisiae* strains in the application environment.

| PEA-producing |  |  | BCHA-producing |  |  |
| --- | --- | --- | --- | --- | --- |
| Selection env. | RS | TS | Selection env. | RS | TS |
| Asp, Met, Trp | $1.93 \times 10^{-2}$ | -0.671 | Arg, Asp, Lys | $2.65 \times 10^{-2}$ | -0.690 |
| Gal, Arg, Gly | $1.88 \times 10^{-2}$ | -0.680 | Arg, NH<sub>4</sub>, Ser | $2.99 \times 10^{-2}$ | -0.695 |
| Gal, Asn, Lys | $1.90 \times 10^{-2}$ | -0.661 | EtOH, Glc, Ser | $2.85 \times 10^{-2}$ | -0.815 |
| Gal, Gol, Asn | $2.24 \times 10^{-2}$ | -0.616 | Gal, Glc, Ile | $5.94 \times 10^{-2}$ | -0.831 |
| Gal, Lys, Phe | $6.61 \times 10^{-2}$ | -0.841 | Gal, Ile, Lys | $4.61 \times 10^{-2}$ | -0.865 |
| Glc, Met, Phe | $6.77 \times 10^{-2}$ | -0.831 | Glc, Ile, Val | $5.69 \times 10^{-2}$ | -0.843 |
| Gol, Asp, Ile | $2.03 \times 10^{-2}$ | -0.640 | Glc, Leu, NH<sub>4</sub> | $5.41 \times 10^{-2}$ | -0.836 |
| Gol, Asp, Trp | $1.79 \times 10^{-2}$ | -0.671 | Glc, Leu, Trp | $6.04 \times 10^{-2}$ | -0.818 |
| Gol, Lys, NH<sub>4</sub> | $1.81 \times 10^{-2}$ | -0.699 | Gol, Ala, Phe | $2.64 \times 10^{-2}$ | -0.660 |
| Gol, Met, Phe | $6.19 \times 10^{-2}$ | -0.854 | Gol, Leu, Met | $5.20 \times 10^{-2}$ | -0.838 |

Gal: D-galactose, Glc: D-glucose, Gol: glycerol, EtOH: ethanol, NH<sub>4</sub>: ammonium, and standard three-letter symbols of amino acids.

**Fig 3.**
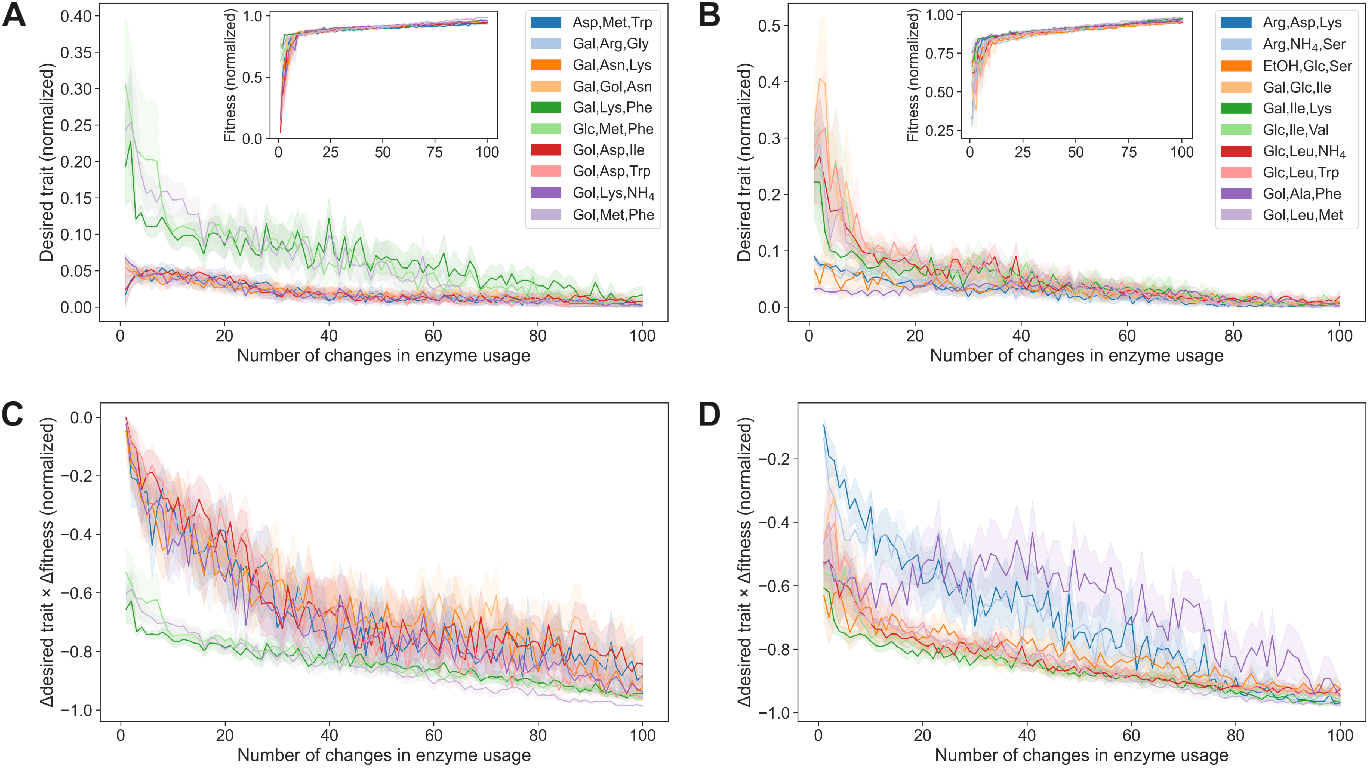
Evolutionary trajectories of ALE-evolved *S. cerevisiae* strains in the application environment. Evolutionary trajectories against the number of changes in enzyme usage (*k* = 1 to 100) for strains evolved in example selection environments for the PEA- and BCHA-producting traits, respectively. Each point in a trajectory represents the mean of up to 30 samples, except that at *k* = 1 all feasible samples are included; shaded bands denote 95% confidence intervals; legend indicates the nutrient compositions of the selection environments and the corresponding trajectory colors. (A, C) PEA-producing trait: (A) normalized desired trait vs. *k* (inset: normalized fitness vs. *k*); (C) normalized Δ(desired trait) × Δ(fitness) vs. *k*. (B, D) BCHA-producing trait: (B) normalized desired trait vs. *k* (inset: normalized fitness vs. *k*); (D) normalized Δ(desired trait)×Δ(fitness) vs. *k*.

Both PEA- and BCHA-producing traits declined with increasing changes in enzyme usage as fitness improved in the application environment (Figure 3A, B), accompanied by increasingly negative trait-fitness trade-off (Figure 3C, D). For the PEA-producing trait, the strains evolved in selection environments containing L-phenylalanine (e.g., `Gal,Lys,Phe`; `Glc,Met,Phe`; `Gol,Met,Phe`) maintained substantially higher trait levels throughout the trajectories (Figure 3A), resulting in markedly higher RS (Table 1). However, these strains also exhibited among the lowest (i.e., most negative) TS (Figure 3C, Table 1), indicating a stronger trade-off with fitness. Similarly, for the BCHA-producing trait, the strains evolved in selection environments containing branched-chain amino acids (e.g., `Gal,Glc,Ile`; `Glc,Ile,Val`; `Glc,Leu,Trp`) tended to retain higher trait levels (Figure 3B) and thus higher RS (Table 1), but at the expense of lower TS (Figure 3D, Table 1). However, this negative association was weaker than for the PEA-producing trait, for instance, TS of `Glc,Leu,Trp` was moderate among the examples.

Overall, the rankings of RS and TS were largely opposite across examples, highlighting a tension between maintaining trait robustness and weakening its coupling to fitness. Among the PEA-producing examples, the strains evolved in selection environments not containing L-phenylalanine showed a relatively low RS but higher TS (Figure 3A and C, Table 1). Likewise, for the BCHA-producing examples, `Arg,Asp,Gln` and `EtOH,Arg,Asn` had markedly lower RS but higher TS (Figure 3B and D, Table 1).

These cases of higher TS indicate a weaker negative coupling between fitness and the desired trait, making such strains promising candidates for further development to achieve higher robustness.

We further examined the dependence of RS and TS on the nutrient composition of the selection environments by fitting a statistical model to estimate the effects of nutrient pairs. Each selection environment contained three nutrients, and nutrient pairs were defined as all three pairwise combinations of these nutrients. As shown in Figure 4A and B, each dot corresponds to one nutrient pair, with its position indicating whether the pair increased or decreased RS (horizontal axis) and TS (vertical axis), and dot size reflecting the number of the selection environments that contained this pair. For both PEA- and BCHA-producing traits, most nutrient pairs lay in the upper-left or lower-right quadrants, showing opposite effects on RS and TS, i.e, they tended to increase RS while decreasing TS, or vice versa. This pattern indicates that strains evolved under a majority of nutrient combinations experience a tension between maintaining trait robustness and weakening its trade-off to fitness. Nevertheless, a few of nutrient pairs still fell in the upper-right quadrant and, thus, showed positive effects on both RS and TS, such as `Glc,Phe` for the PEA-producing trait. Consistent with these observations, the effect coefficients on RS and TS were moderately negatively correlated for both traits. These findings suggested that, although ALE-evolved strains exhibiting both high RS and high TS were generally rare, certain nutrient combinations could partially alleviate this tension.

**Fig 4.**
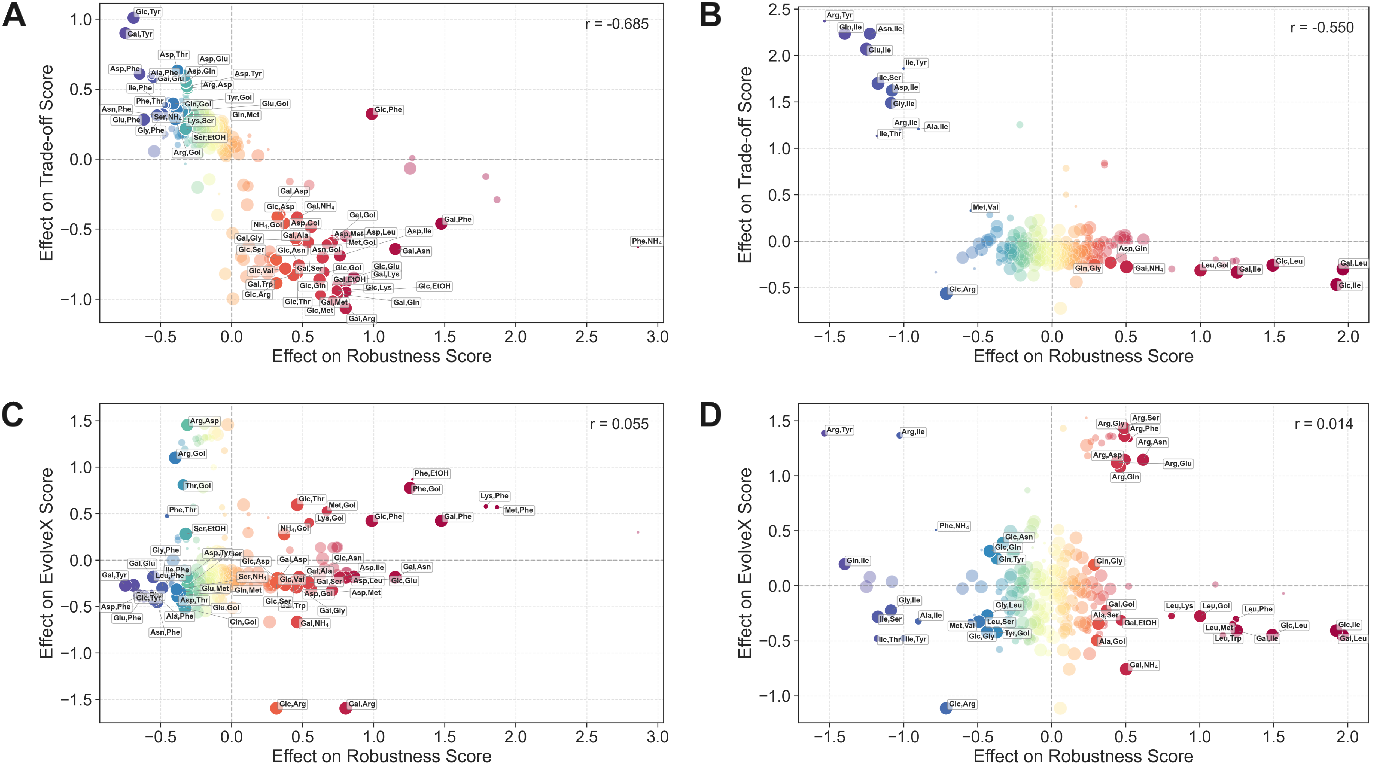
Nutrient pair effects on Robustness Score (RS), Trade-off Score (TS), and EvolveX Score. Each dot is a nutrient pair; axes show the effect coefficients of that pair on the indicated score. Dots with coefficient *p*-value *<* 0.05 are highlighted and annotated. (A, B) Pearson correlation across effect on RS vs. effect on TS for (A) PEA-producing: *r* = −0.685 (*p <* 0.05); (B) BCHA-producing: *r* = −0.550 (*p <* 0.05). (C, D) Pearson correlation across effect on RS vs. effect on EvolveX Score for (C) PEA-producing: *r* = −0.055 (*p* ≫0.05); (D) BCHA-producing: *r* = 0.014 (*p* ≫0.05). Dot size reflects the number of the selection environments containing the nutrient pair. Dashed lines mark zero effect. *r* values are shown in each panel.

We also applied the same statistical analysis to compare RS with the EvolveX Scores reported in [27]. As shown in Figure 4C and D, the nutrient pair effects on the EvolveX Scores were very weakly correlated with those on RS, but the nutrient pairs were clearly separated into two halves along the RS axis. Notably, for the PEA-producing trait, some L-phenylalanine-containing nutrient pairs, such as `Lys,Phe` and `Met,Phe`, showed positive effects on both RS and the EvolveX Score (Figure 4C); for the BCHA-producing trait, some arginine-containing nutrient pairs, such as `Arg,Ser` and `Arg,Glu`, showed similar positive effects (Figure 4D), suggesting that these combinations favor both enhanced trait level and robustness in the application environment.

## Discussion

Developing strains that robustly maintain desired trait expression in industrial applications remains a major challenge in microbial biotechnology. Here, we addressed this challenge with the computational framework TRAEV that can produce predictions aligned with experimental observations, and thus, complement the existing model-guided strain design methods such as EvolveX [27] for non-GMO strain development or growth-product coupling by genetic engineering [33]. This would alleviate the immense resource need for the proposed experimental stability screening of strains [10].

Prior to our work, several methods have previously been proposed for predicting and quantifying biological robustness [34–37]. They share a common feature that the robustness is evaluated with respect to a predefined set of perturbations in a largely static system rather than simulating evolutionary trajectories. In contrast, TRAEV explicitly predicts trajectories of evolutionary trait adaptations. Thus, the robustness is predicted not only as sensitivity to perturbations but also in terms of how traits and fitness co-evolve via sub-optimally adapted intermediate states in the application environment. This trajectory-based perspective is particularly relevant for industrial processes, where microbial strains undergo notable biomass expansion to production scale and potentially prolonged cultivation.

The two case studies illustrated proposed applications of TRAEV for designing strategies for robust industrial strain development with biological explainability. In the pigment production case, the relative trait robustness in *S. cerevisiae* predicted using TRAEV was consistent with the experimental observations [31], indicating that TRAEV can simulate key adaptive changes in metabolic pathways during a long-term cultivation. At the same time, the explainability of GEM simulations provided a hypothesis of the mechanisms underlying the observed difference in robustness. Indigoidine is synthesized from L-glutamine [11], which is directly linked to both the citric acid cycle and nitrogen metabolism, making its synthesis sensitive to many enzyme-usage changes in central metabolism when nutrient utilization is already efficient. In contrast, the bikaverin-synthesizing *S. cerevisiae* S288C strain can gain improved fitness via many enzyme-usage changes, including those enhancing galactose utilization [31], that do not impair bikaverin synthesis from acetyl-CoA through a relatively insulated pathway [12].

In the aroma production case, TRAEV was used prospectively to evaluate and prioritize selection environments for developing robust non-GMO aroma producing *S. cerevisiae* strains via ALE. The results showed that the robust PEA-producing trait could be developed using a narrower spectrum of selection environments, primarily those containing L-phenylalanine, than the BCHA-producing trait. Both PEA (phenylethyl alcohol and its acetate ester) and BCHA (branched-chain amino acid derived aromas) are generated via the Ehrlich pathway [38] but L-phenylalanine, the precursor of PEA, is more expensive for yeast to synthesize (e.g., in terms of NADPH requirement) than the branched-chain amino acids [39, 40]. In addition to the higher requirement for the supply of reducing power, L-phenylalanine synthesis is more connected to other central pathways (i.e., glycolysis, pentose-phosphate pathway, and nucleotide synthesis) than branched-chain amino acid synthesis. Furthermore, the L-phenylalanine metabolism via Ehrlich pathway involves a specific decarboxylating enzyme (ARO10) while branched-chain amino acid metabolism after deamination occurs by promiscuous enzymes.

The nutrient pair effect analysis further showed that the combinations rarely exerted strong effects on both high trait robustness and less negative trait–fitness trade-off, consistent with the traits and fitness often exhibiting inherent trade-offs due to limited cellular resources [41, 42]. Nevertheless, some nutrient combinations were predicted to select for strains with relatively high RS and less negative TS, partially alleviating the typical trade-off between traits and fitness. The analysis provided also complementary insights to EvolveX Scores [27]. Nutrient combinations having similar EvolveX Scores were further stratified into subsets with higher and lower predicted robustness. Thus, TRAEV could refine chemical environments suitable for adaptively evolving desired traits designed using EvolveX by ranking them according to predicted trait robustness and trait–fitness trade-off of the developed strain in the application environment.

Despite the advantages, the current TRAEV implementation is limited by the models it incorporates. First, the modeled application environment did not account for temporal changes during cell growth, such as sequential nutrient depletion or increasing ethanol concentration in a wine fermentation. Modeling time-varying environments (e.g., using dynamic FBA [43]) would allow robustness to be assessed under realistic conditions but requires data on compound utilization and production dynamics. Second, incorporating models of trait-fitness dependencies extended beyond metabolism would improve simulation fidelity and predictive accuracy as metabolism is not isolated from other cellular processes.

Ultimately, we envision TRAEV could become a practical component of strain and process development workflows and contribute to enhancing the economic attractiveness of industrial applications using microbial cells.

## Methods

### Data, models, and software

The wild-type genome-scale metabolic model (GEM) used in this study was the yeast consensus GEM (v9.0.2, https://zenodo.org/records/14210050) [32], transformed using the Gene-Protein-Reaction (GPR) formulation [44] implemented in the `ReFramed` package (v1.5.1, https://github.com/cdanielmachado/reframed). In contrast to the basic GEM, the GPR-transformed model introduces variables representing enzyme usages and distinguishes among isoenzymes catalyzing identical metabolic reactions. The `ReFramed` package relies on the `COBRApy` package (v0.29.1) [45] for metabolic modeling and on the Gurobi Optimizer (v11.0.1) for solving optimization problems.

### Definition of fitness and desired trait

The biomass yield was used as a proxy for the fitness *f* of a strain in metabolic state **v**, defined as:

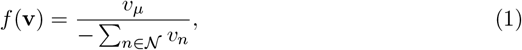

where *v*_*µ*_ denotes the specific growth rate (h^−1^), and 𝒩 represents the set of nutrient uptake reactions. For each reaction *n* ∈ 𝒩, *v*_*n*_ denotes its flux (mmol · gDW^−1^ · h^−1^), which is non-positive by the convention of directionality. Thus, the denominator is a positive value representing the total nutrient uptake.

The desired trait *d* of a strain in metabolic state **v** was defined as the yield of the target product(s):

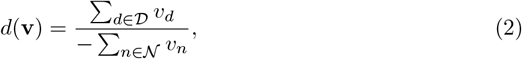

where 𝒟 denotes the set of secretion reactions of the target product(s), and *v*_*d*_ represents the flux of each reaction *d* ∈ 𝒟. The denominator is the same as that in Eq. 1. The reactions included in 𝒟 for the experiments in this study are listed in S2 File.

The fitness and desired trait values were normalized relative to their respective baselines as follows:

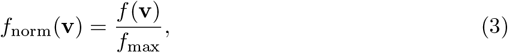

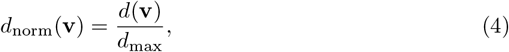

where *f*_max_ and *d*_max_ denote the normalization baselines. Specifically, *f*_max_ is the fitness value of the strain at **v**_a_. *d*_max_ is the trait value of the strain at **v**_p_ when comparing different desired traits. However, when comparing the level of the same desired trait across multiple strains, *d*_max_ is instead set to the maximum trait value among all strains.

### Calculation of v_r_

#### pFBA (FBA)

The metabolic state of the reference strain (**v**_r_) for TRAEV was calculated using pFBA [29]. pFBA is an extension of FBA [22], in which a second optimization step that minimizes the total flux is performed after solving the FBA problem, reflecting the principle that cells tend to economize resources.

For simulating adaptive laboratory evolution (ALE), the FBA problem was formulated as follows:

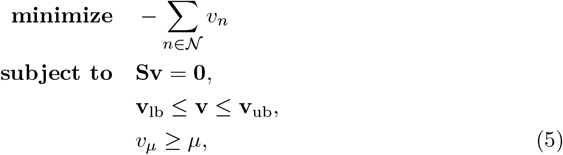

where **S** is the stoichiometric matrix of the GEM, and **v** represents the metabolic state, i.e., the vector (distribution) of both metabolic fluxes and enzyme usages. **v**_lb_ and **v**_ub_ denote the lower and upper bounds of **v**, respectively, and the specific growth rate *v*_*µ*_ is constrained to be at least *µ*.

Since using pFBA in this study, after solving Eq. 5 to obtain the optimal objective value (i.e., minimum nutrient uptake), a secondary optimization was performed to minimize the total flux, subject to maintaining the FBA-derived optimal objective.

For simulating genetic modification (GM), the objective in FBA (Eq. 5) was modified to maximization of ∑_*d* ∈𝒟_ *v*_*d*_, and the lower bounds of *v*_*n*_ were constrained to physiologically feasible values.

#### SWITCHX

In addition to pFBA, we used an alternative method, termed SWITCHX, for cases where the metabolic state of the wild type (parental) strain is available. SWITCHX implementation derives from regulatory on/off minimization (ROOM) [46]. It minimizes the total number of flux and/or enzyme usage changes (switches) relative to the wild type or parental state, while additionally imposing a constraint that fixes the nutrient uptake to the minimum value obtained from FBA (Eq. 5) to ensure physiological optimality.

For simulating both ALE and GM, given the metabolic state **w** of the wild-type (parental) strain, the SWITCHX problem was formulated as follows:

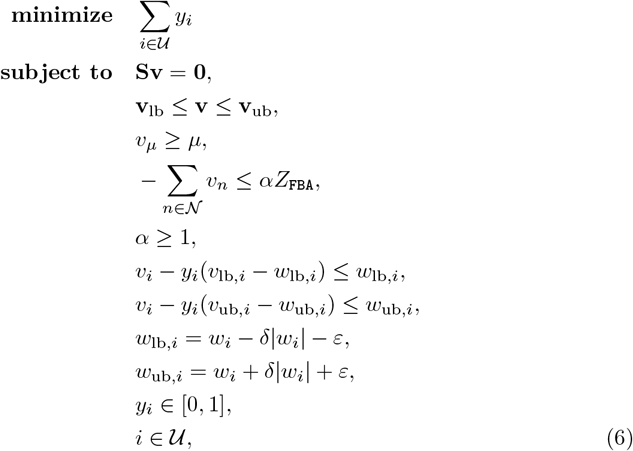

where **S** is the stoichiometric matrix of the GEM, and **v** represents the metabolic state, i.e., the vector (distribution) of both metabolic fluxes and enzyme usages. **v**_lb_ and **v**_ub_ denote the lower and upper bounds of **v**, respectively, and the specific growth rate *v*_*µ*_ is constrained to be at least *µ*. The total nutrient uptake is limited by *αZ*_FBA_, where *Z*_FBA_ denotes the minimum total nutrient uptake obtained from FBA (Eq. 5), and *α* is a scaling parameter controlling the allowed deviation from *Z*_FBA_. In this study, *α* was fixed at 1 to ensure achieving *Z*_FBA_. *y*_*i*_ is a binary variable representing whether the value of *v*_*i*_ (*i*-th component of **v**) deviates from the wild-type range [*w*_lb,*i*_, *w*_ub,*i*_]. The bounds *w*_ub,*i*_ and *w*_lb,*i*_ are derived from the corresponding *w*_*i*_ (*i*-th component of **w**) using the parameters *δ* and *ε*, as in ROOM [46]. 𝒰 denotes the set of all components involved, which in this study are all enzyme usages.

#### TRAEV

We proposed the TRAEV framework to perform the Simulation and Robustness Analysis of the adaptive evolution of developed strains in application environments. In the Simulation, we calculated the strain’s immediate phenotypically adjusted metabolic state (**v**_p_) and evolutionarily adapted metabolic state (**v**_a_) in the application environment.

#### Calculation of v_p_

The calculation of **v**_p_ was implemented using linear MOMA [30], a linear version of quadratic MOMA [23], with **v**_r_ as the reference state. MOMA assumes that cells try to minimize deviation from their previous metabolic state when a sudden change (e.g., environmental shift) occurs. This reflects the short-term, physiological plastic response before evolutionary adaptation [28].

The objective of quadratic MOMA is to minimize the Euclidean (L2-norm) distance to the reference state, denoted as **w**, and the problem is formulated as follows:

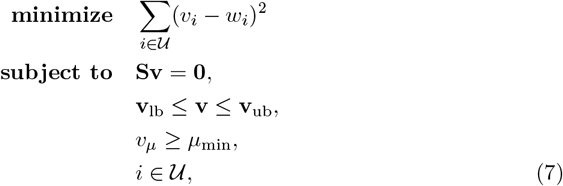

where **S** is the stoichiometric matrix of the GEM, and **v** represents the metabolic state, i.e., the vector (distribution) of both metabolic fluxes and enzyme usages. **v**_lb_ and **v**_ub_ denote the lower and upper bounds of **v**, respectively, and the growth rate *v*_*µ*_ was constrained to be at least *µ*_min_, a biologically minimum growth rate. *v*_*i*_ denotes the *i*-th component of **v** and *w*_*i*_ denotes the *i*-th component of **w**. 𝒰 denotes the set of all components involved, which in this study are all enzyme usages.

In linear MOMA, the objective function in Eq. 7 is replaced with the L1-norm distance ∑_*i* ∈𝒰_| *v*_*i*_ − *w*_*i*_ |. The quadratic MOMA tends to produce a solution in which all of the components deviate slightly from the reference state. In contrast, linear MOMA tends to produce a solution in which most of the components are the same as the reference, while a few deviating significantly (a typical effect of the L2 norm versus the L1 norm) [47]. This feature can make the solutions more biologically interpretable.

#### Calculation of v_a_

The calculation **v**_a_ was implemented using a greedy algorithm 1 built on SWITCHX (Eq. 6).

##### Algorithm 1

Greedy search algorithm for calculating **v**_a_

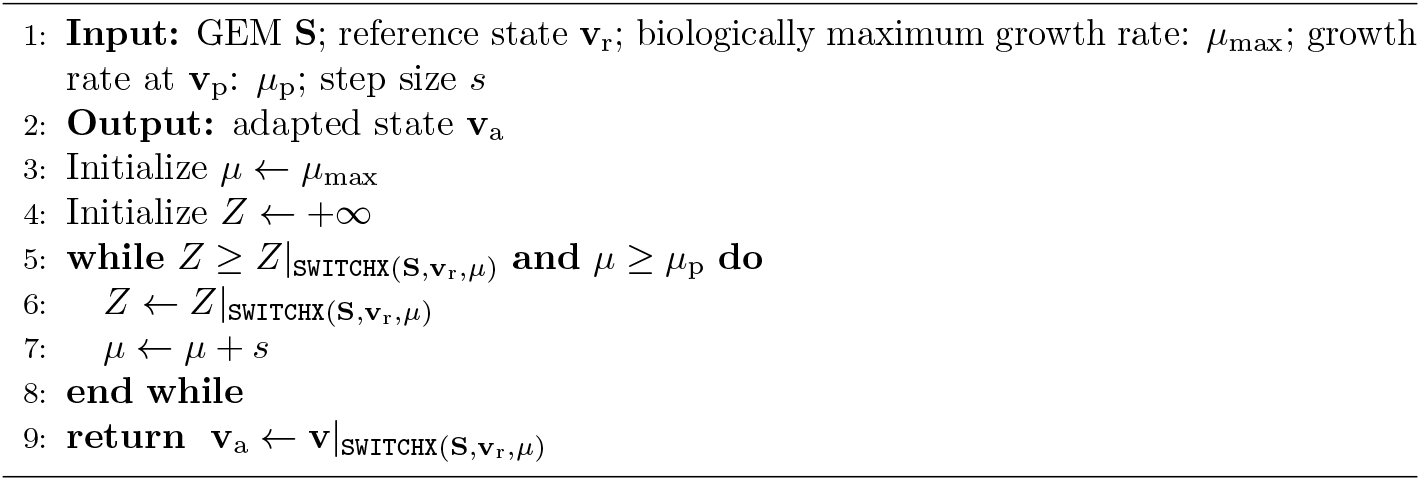

Since FBA is integrated into SWITCHX, the solution obtained for any specified value of the growth rate (*µ*) always achieves the optimal fitness (Eq. 1). To estimate the evolutionarily adapted value of *µ* and the corresponding **v**_a_, we developed a greedy search algorithm (Algorithm 1). Starting from a biologically reasonable maximum growth rate (*µ*_max_), the algorithm decreases *µ* stepwise, resolves SWITCHX at each step, and stops at the first local minimum of the number of changes in enzyme usage relative to the reference state **v**_r_ (i.e., the SWITCHX objective value *Z*).

The algorithm is built on the assumption that evolution tends to achieve the maximum possible growth rate while incurring the least changes in enzyme usages (i.e., mutations). Thus, the algorithm is a simple one-dimensional greedy heuristic that preferentially selects a high growth, minimally perturbed state rather than guaranteeing the global minimum of *Z* over all feasible *µ*. Despite lack of global optimality, the method is computationally efficient, easy to interpret, and provides a practical, evolutionarily grounded approximation of the adapted growth rate in our setting.

#### Robustness Analysis

Following the Simulation, the Robustness Analysis was performed through a Monte Carlo-based perturbation analysis method.

Let ℳ denote the set of perturbations (i.e., enzyme usages that change) between **v**_p_ and **v**_a_, and let |ℳ| be its cardinality. For each perturbation count *k* ∈ ·1, 2, …, |ℳ |}, there are 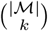 possible perturbation combinations. From these %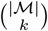 perturbation combinations, we randomly selected a sample *s*, which is a subset of ℳ and its corresponding evolutionarily intermediate metabolic state was defined as follows:

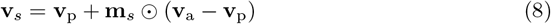

where **m**_*s*_ is a binary mask whose component is 1 if the corresponding perturbation is included in *s* and 0 otherwise. To guarantee the feasibility of **v**_*s*_, we did not use Eq. 8 directly. Instead, quadratic MOMA (Eq. 7) was run with **v**_p_ as the reference state and an additional constraint that the perturbations in *s* were fixed to their values in **v**_a_. If MOMA returned a feasible solution, it was taken as the feasible approximation of **v**_*s*_, where the components in *s* equal to the values in **v**_a_, whereas the remaining components are kept close to their values in **v**_p_. We repeated the above sampling procedure for *k* from 1 up to *m* ≤ |ℳ |, and for each *k* we collected up to *n* feasible evolutionarily intermediate metabolic states. Thus, we obtained a sample set 𝒮 of up to *m* × *n* evolutionarily intermediate metabolic states between **v**_p_ and **v**_a_.

Next, the fitness (Eq. 1) and the desired trait (Eq. 2) values were derived from 𝒮 and normalized (Eqs. 3, 4) for the prediction of the evolutionary trajectories. Specifically, for each *s* ∈ 𝒮, we plotted the normalized fitness *f*_norm_(**v**_*s*_), the normalized desired trait *d*_norm_(**v**_*s*_), and Δ*d*_norm_(**v**_*s*_)Δ*f*_norm_(**v**_*s*_)^1^, against the perturbation count (i.e., number of changes in enzyme usage) of *s*.

Finally, the Robustness Score (RS) and the Trade-off Score (TS) were calculated as follows:

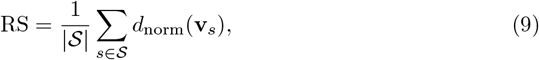

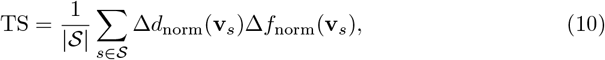

where |𝒮| denotes the cardinality of 𝒮.

### Experimental procedures

#### Simulations for predicting the robustness of heterologous pigment production in *S. cerevisiae*

We augmented the yeast consensus GEM (v9.0.2) [32] for two engineered *S. cerevisiae* strains (the indigoidine-producing CEN.PK113-7D and the bikaverin-producing S288C) with the corresponding heterologous reactions, respectively (see S1 File).

For the indigoidine-producing strain, the indigoidine exchange reaction was defined as the desired trait (see S2 File). The environment was set under aerobic conditions with D-glucose as the sole carbon source and ammonium as the sole nitrogen source. A minimum growth rate of 0.3 h^−1^ and a maximum D-glucose uptake rate of ≤10 mmol · gDW^−1^ · h^−1^ were enforced, while ammonium uptake was left unconstrained (≤1000mmol · gDW^−1^ · h^−1^). pFBA was then performed with the objective of maximizing the indigoidine secretion subject to these constraints.

Analogously, for the bikaverin-producing strain, the bikaverin exchange reaction was defined as the desired trait (see S2 File). The environment was set under aerobic conditions with D-glucose as the sole carbon source and ammonium as the sole nitrogen source. A minimum growth rate of 0.3 h^−1^ and a maximum D-glucose uptake rate of 10 mmol · gDW^−1^ · h^−1^ were enforced, while ammonium uptake was left unconstrained. pFBA was then performed with the objective of maximizing the bikaverin secretion subject to these constraints.

We next applied TRAEV to both strains (indigoidine-producing and bikaverin-producing) in the two application environments: YNBG (ammonium + D-galactose) and YPG (peptone + D-galactose), respectively. For the YNBG simulations, the uptake of D-galactose and ammonium was set to be unconstrained; for the YPG simulations, the D-galactose uptake was unconstrained, whereas the uptake of the 20 standard amino acids was permitted but capped at 1 mmol · gDW^−1^ · h^−1^. The maximum growth rates (*µ*_max_) were set to 0.4 h^−1^ for YNBG and 0.6 h^−1^ for YPG, referring to [48]. In the subsequent Robustness Analysis, we set the maximum perturbation count *m* = 100 and the maximum sample size *n* = 30. Finally, the TRAEV outputs were exported for the prediction of the evolutionary trajectories and the calculation of RS.

#### Simulations for predicting the robustness of aroma production in evolved *S. cerevisiae* strains under different selection environments

We adjusted the yeast consensus GEM (v9.0.2) [32] to favor aroma compound production (see S1 File). To simulate the aerobic ALE process in each selection environment, the uptake fluxes of nutrients present in the environment were set to unconstrained (≤ 1000mmol · gDW^−1^ · h^−1^), while the uptake fluxes of nutrients absent from the environment were set to zero. The growth rate *µ* = 0.4 h^−1^ was enforced. pFBA was then performed with the objective of minimizing total nutrient uptake. Simulations were conducted across all 1171 selection environments (see S3 File), resulting in 1171 reference strains.

For each reference strain, we applied TRAEV to simulate adaptive evolution in an anaerobic wine must-like environment (see S1 File). The adapted growth rate was calculated using Algorithm 1. The desired traits related to PEA and BCHA production were calculated as the sum of fluxes through the aroma exchange reactions (listed in S2 File). Next, the Robustness Analysis was executed with the maximum perturbation count *m* = 100 and the maximum sample size *n* = 30. Finally, the results were exported for the prediction of the evolutionary trajectories, calculation of RS and TS, and further statistical analysis.

We ran the program across all 1,171 selection environments for the PEA and BCHA production traits in parallel on the Aalto Triton cluster using ≈ 100 CPU cores, completing in ≈ 3 days of wall-clock time.

### Statistical analysis

Statistical correlation analyses were conducted to examine the associations between nutrient pair effects across different scores. Because each selection environment contained three nutrients, many nutrient pairs recurred in multiple environments, enabling statistical analyses and yielding more informative insights than the original scores.

For each selection environment, each score (i.e., RS, TS, or EvolveX Score) corresponding to that environment was standardized using Z-score normalization with the mean and standard deviation computed across all the selection environments. The resulting standardized score was treated as the response variable, denoted as *y*. The presence of nutrient pair *i* across all environments was encoded as a binary predictor variable, denoted as *x*_*i*_. For all *P* nutrient pairs, we modeled the association between *x*_*i*_ and *y* using ordinary least squares (OLS) regression as follows:

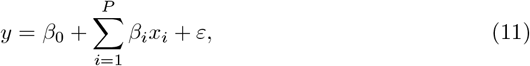

where *β*_*i*_ is the estimated coefficient representing the effect of nutrient pair *i* on the score, and *ε* is the error term. The analysis was implemented using the `statsmodels` package (v0.14.5). For each estimated coefficient, a corresponding *p*-value was computed.

After fitting the OLS regression models respectively for RS, TS, and EvolveX Score and obtaining the corresponding effect coefficients and associated *p*-values, Pearson correlation coefficients were calculated between the sets of effect coefficients from the two pairs of scores (RS vs. TS, and RS vs. EvolveX Score) across all *P* nutrient pairs. Because the original EvolveX Scores indicated better performance with lower values, they were reversed prior to all the analyses to ensure directional consistency with the other scores, such that higher values consistently reflected better performance.

## Code and data availability

All code and the corresponding data (including results) are available at https://zenodo.org/records/18097289.

## Supporting information

**S1 File. Supplementary information**. Additional modeling information for strains and environments.

**S2 File. Desired traits**. List of metabolic reactions representing the desired trait of strains.

**S3 File. Selection environments**. List of selection environments for ALE of aroma-producing strains.

## Footnotes

1 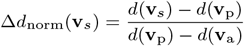 and 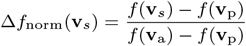

## Notes

https://zenodo.org/records/18097289

### Competing Interest Statement

The authors have declared no competing interest.

### Funding

- Research Council of Finland: PJ: 329930 and 352417, VM: 329931
- Novo Nordisk Foundation: PJ: NNF22OC0080180

