## Supplementary material for "*In silico* prediction of metabolic trait robustness in microbial cells": S1

### S1.1 GEM modifications

The wild-type genome-scale metabolic model (GEM) used in this study was the yeast consensus model (v9.0.2, `https://zenodo.org/records/14210050`) [1]. In addition, the reactions (2R,3S)-3-methylmalate:NAD$^+$ oxidoreductase (`r_4581`) and malate/$\beta$-methylmalate synthase (`r_4582`) were closed. Both reactions are annotated with Confidence Level 0 and lack gene associations; therefore, closing them can avoid spurious side routes and keep flux in well-supported metabolic pathways.

#### S1.1.1 Indigoidine-producing strain

The GEM for an indigoidine-producing strain was constructed by incorporating the indigoidine biosynthetic pathway into the yeast consensus model. The pathway comprises a heterologous gene encoding indigoidine synthetase (BpsA) and a phosphopantetheinyl transferase (PPTase). BpsA is a nonribosomal peptide synthetase (NRPS) that catalyzes the condensation of two L-glutamine molecules to form one indigoidine molecule, while PPTase activates BpsA by converting it from its apo- to holo-form [2]. Because the PPTase reaction is a post-translational modification that does not influence metabolic fluxes, it was excluded from the model. Accordingly, the indigoidine biosynthetic pathway was represented as a single overall reaction based on the stoichiometry described in [3], omitting the FMN-dependent step since the flavin cofactor functions as an enzyme-bound prosthetic group and therefore does not carry metabolic flux. The added synthesis, transport, and exchange reactions are listed in Table S1.

**Table S1. Reactions added for the indigoidine biosynthetic pathway.**

| No. | Reaction Name | Stoichiometry | Compartment | Gene Association |
|---|---|---|---|---|
| 1 | Indigoidine synthase | 2L-glutamine + 2ATP + $O_2$ $\rightarrow$ Indigoidine[c] + 2AMP + $2PP_i$ + $2H_2O$ | cytosol | *bpsA* |
| 2 | Indigoidine transport | Indigoidine[c] $\rightleftharpoons$ Indigoidine[e] | cytosol $\rightleftharpoons$ extracellular | – |
| 3 | Indigoidine exchange | Indigoidine[e] $\rightarrow$ | extracellular | – |

#### S1.1.2 Bikaverin-producing strain

The GEM for an bikaverin-producing strain was constructed by incorporating the bikaverin biosynthetic pathway into the yeast consensus model. The pathway essentially comprises three heterologous genes encoding one polyketide synthase (Bik1) and two tailoring enzymes (Bik2, Bik3), and a phosphopantetheinyl transferase (PPTase). Bik1, activated by PPTase, condenses one acetyl-CoA and eight malonyl-CoA units to form pre-bikaverin, which is subsequently oxidized and methylated by Bik2 (monooxygenase) and Bik3 (O-methyltransferase) to produce bikaverin [4]. Thus, the bikaverin biosynthesis was represented as a single overall reaction catalyzed by Bik1-3. The added synthesis, transport, and exchange reactions are listed in Table S2.

**Table S2. Reactions added for the bikaverin biosynthetic pathway.**

| No. | Reaction name | Stoichiometry | Compartment | Gene association |
|---|---|---|---|---|
| 1 | Bikaverin synthase | 8Malonyl-CoA + Acetyl-CoA + 2SAM + $2O_2$ + 2NADPH$\rightarrow$ Bikaverin[c] + 9CoA + $8CO_2$ + 2SAH + $2H_2O$ + $2NADP^+$ | cytosol | *bik1*, *bik2*, and *bik3* |
| 2 | Bikaverin transport | Bikaveri[c] $\rightleftharpoons$ Bikaverin[e] | cytosol $\rightleftharpoons$ extracellular | – |
| 3 | Bikaverin exchange | Bikaverin[e] $\rightarrow$ | extracellular | – |

#### S1.1.3 Aroma-producing strain

For the aroma-producing strains, both the phenylethyl alcohol (PEA) and branched-chain higher alcohol (BCHA) pathways were adjusted in the GEM to favor aroma compound synthesis while remaining physiologically reasonable. Specifically, the reactions isoamyl acetate-hydrolyzing esterase (`r_0656`) and isobutyl acetate-hydrolyzing esterase (`r_0657`) were closed. These constraints suppress the hydrolysis of higher alcohol acetate esters, eliminate futile ester cycles, and enable the model to reproduce the increased accumulation of acetate esters.

Additionally, the phenylacetaldehyde secretion (`r_2001`) was closed. Phenylacetaldehyde is an intermediate of the Ehrlich pathway that is typically reduced intracellularly to PEA rather than secreted. Closing its secretion in the GEM redirects the flux toward PEA formation.

#### S1.1.4 Wine must environment

The wine must environment represents the nutrient and physicochemical conditions that yeast *Saccharomyces cerevisiae* encounters during grape must fermentation. This environment is largely (after early phase) anaerobic, with D-glucose and D-fructose serving as the primary carbon sources and yeast assimilable nitrogen (ammonium and free amino acids) as the main nitrogen sources. Under these conditions, yeast metabolism relies predominantly on fermentation to produce ethanol, glycerol, and aroma compounds, while oxidative respiration remains inactive.

To simulate this environment *in silico*, the GEM was configured to allow unconstrained uptake ($\leq 1000 \mathrm{mmol} \cdot \mathrm{gDW}^{-1} \cdot \mathrm{h}^{-1}$) of D-glucose (`r_1714`), D-fructose (`r_1709`), and ammonium (`r_1654`), while capping uptake of each of the 20 standard amino acids (`r_1873`, `r_1879`, `r_1880`, `r_1881`, `r_1883`, `r_1889`, `r_1891`, `r_1810`, `r_1893`, `r_1897`, `r_1899`, `r_1900`, `r_1902`, `r_1903`, `r_1904`, `r_1906`, `r_1911`, `r_1912`, `r_1913`, `r_1914`) at $1\ \mathrm{mmol} \cdot \mathrm{gDW}^{-1} \cdot \mathrm{h}^{-1}$ to prevent them from serving as major carbon sources. Anaerobiosis was mimicked by closing reactions associated with oxidative respiration, including cytochrome c oxidase (`r_0438`), succinate dehydrogenase (`r_1021`), mitochondrial ATP synthase (`r_0226`), and FMN reductase (`r_0441`, `r_0441`).

Furthermore, reactions involved in glycerol catabolism (`r_0487`, `r_0488`) and glyoxylate cycle activity (`r_0662`) were closed to ensure that intracellularly produced glycerol serves primarily to maintain redox balance rather than be fed back into central carbon metabolism. This is consistent with experimental evidence that glycerol accumulates and is secreted during anaerobic fermentation.

In addition, isocitrate dehydrogenase (`r_0659`), glycine hydroxymethyltransferase (`r_0502`, `r_0503`), and methylenetetrahydrofolate dehydrogenase (`r_0732`, `r_0733`), were constrained to proceed only in their physiologically forward directions. Because the reverse directions of these reactions are thermodynamically unfavorable under fermentative conditions, applying directional constraints ensures physiologically realistic NADH/NADPH cofactor balances.
